# Foam rolling intervention improves lactate clearance after high-intensity exercise

**DOI:** 10.1101/2024.02.16.580741

**Authors:** Kazuki Kasahara, Keita Oneyama, Takeru Ito, Masatoshi Nakamura, Genta Ochi

**Affiliations:** Sanyudo hospital, Fukuda Town, Yonezawa City, Yamagata, Japan; Department of Health and Sports, Niigata University of Health and Welfare, Niigata, Japan; Faculty of Rehabilitation Sciences, Nishi Kyushu University, 4490-9 Ozaki, Kanzaki, Saga, 842-8585, Japan; Institute for Human Movement and Medical Sciences, Niigata University of Health and Welfare, Niigata, Japan

**Keywords:** foam roller, fatigue, Profile of Mood States 2nd Edition (POMS2), recovery, cognitive function

## Abstract

**Purpose:** The effects foam rolling (FR) intervention on lactate clearance and the impaired executive function associated with fatigue after high-intensity exercise remain unclear. This study examined whether FR is an effective tool for fatigue recovery.

**Methods:** Eighteen healthy adults (10 males, 8 females; 21.8±1.7 years) without consistent exercise habits participated. The participants performed high-intensity exercises using an ergometer with a progressive load. Post-exercise FR intervention was compared with the control condition. Measurements included lactate, vigor/fatigue by the Profile of Mood States 2nd edition, cognitive function (cognitive task performance), and leg and body rating of perceived exertion (RPE) pre-and post-exercise, and post-intervention (5 min post-exercise).

**Results:** The blood lactate concentration post-FR intervention (−7.3±3.0 mmol/L) was significantly reduced. Increased lactate clearance by FR correlated with faster recovery of executive function for those with greater lactate clearance, although cognitive fatigue was not observed after high-intensity exercise (*P* = .086, *r* = 0.41). Lactate clearance was not significantly correlated with RPE in the FR condition, whereas RPE decreased with increased lactate clearance for those with greater lactate clearance in the control condition (leg; *r* = 0.778, body; *r* = 0.669).

**Conclusions:** These results indicated FR intervention may be an effective recovery tool for exhausting exercises.

## INTRODUCTION

Many sports competitions involve high-intensity interval exercises involving repeated sprints and jumps. Ball sports, such as basketball and handball, require high-intensity interval exercises for extended periods with little time between halves or sets. High-intensity interval exercise repeated with short rest periods, results in a noticeable decrease in performance in the second half of matches. This resulting performance loss may include muscle function as well as executive function.^1–3^ Executive function is the foundation of tactical understanding and execution, which includes the ability to make selective decisions during sport activities,^4^ and is a major determinant to athletic performance. Cook and Beaven^5^ reported that recovery to a good psychological state post-exercise is important for maintaining performance, whereas recovery of both peripheral and central functions post-exercise is an urgent issue. However, existing recovery methods, such as cold water immersion^6^ and full-body aerobic exercises, including a rowing ergometer,^7^ are difficult to implement during a game because of the time, location, and competition rules. To maintain mental and physical functions until the latter half of a match, a new recovery method that can be implemented in a limited space and short duration is required.

Muscle fatigue caused by high-intensity exercise is related to acidosis from lactate production.^8^ Therefore, many recovery protocols or methods target lactate clearance.^9,10,11,12^ Interestingly, increased peripheral blood lactate concentration induced by strenuous exercise or intravenous infusion of lactate solution is inversely related to attentional task performance,^13^ suggesting that increased peripheral blood lactate concentration may be a biomarker for cognitive fatigue because lactate crosses the blood-brain barrier. Improving adequate lactate clearance may improve muscle fatigue and executive dysfunction, thereby improving athletic performance; however none of the previous recovery methods have focused on executive function.

Foam rolling (FR) is commonly used as a recovery method. Adamczyk et al.^14^ evaluated the effects of FR intervention after squat jumps at maximal effort for 1 min, which significantly increased lactate clearance for the FR condition than the intervention condition at 30 min post-exercise. In addition, Akinci et al.^15^ implemented circuit-based high-intensity exercise at 85% intensity, as calculated from the heart rate reserve method, and measured lactate levels before exercise training, before the recovery period, and at 5 and 20 min into the recovery period to examine the effects of FR intervention on lactate levels. They reported that FR intervention did not alter lactate concentrations. However, these studies may not have adequately evaluated the effects of these exercises on primarily active and fatigued muscles during exercise because of the long time elapsed after the exercise and it is a whole-body exercise.

This study purported to determine whether FR improves lactate clearance after exercise with fatigue and consequently improve impaired executive function.

## METHODS

### Experimental Approach to the Problem

This study examined the immediate effects of FR, which can be performed in a confined space for a limited period, on lactate clearance and cognitive function after high-intensity exercise with progressive loading. In addition, this study assessed these effects separately in males and females.

A randomized repeated-measures experimental design was used to compare the two conditions: control and FR. Participants were instructed to visit the laboratory twice, with a ≥48 h break. Measurements were conducted before the intervention (pre), after the intervention (post), and 5 min post-intervention. The completed psychological scales included the Profile of Mood States 2nd Edition (POMS2)^16^ and participant ratings of perceived exertion (RPE).^17^ The parameters measured were executive function, psychological scale scores, and blood lactate concentration.

### Participants

A total of 22 right-handed Japanese-speaking young adults (12 males and 10 females) were included in this study conducted between October 2023 and January 2024. The inclusion criteria were: no regular exercise habits; no daily use of a foam roller; native Japanese speakers; and no history of neurological, psychiatric, or respiratory diseases. All participants were instructed to refrain from eating or drinking anything other than mineral water 2 h before starting the experiment. In this study, analyses were conducted with data from 18 participants (10 males and 8 females); four participants were excluded because of their inability to perform the FR intervention after high-intensity exercise. Table 1 presents the participant characteristics. We performed a post-hoc sensitivity analysis using this sample with 80% power and an α = 0.05 demonstrated sufficient sensitivity to detect repeated-measures effects exceeding *f* = 0.31 as computed using G*Power (3.1.9.2; The G*Power Team).

**Table 1.** Participant characteristics (mean±SD)

| Characteristic | Males (N=10) | Females (N=8) |
| --- | --- | --- |
| Height (cm) | 172.1±5.0 | 158.4±5.0* |
| Weight (kg) | 67.1±8.4 | 48.1±5.3* |
| Age (years) | 22.4±1.3 | 21.0±2.0 |
| Thigh circumference (Right) (cm) | 53.1±4.9 | 46.7±6.0* |
| Thigh circumference (Left) (cm) | 53.2±4.3 | 47.1±5.9* |

\*: Significant differences were found for females compared to males ( $P < .05$ ). SD, standard deviation.

This study was conducted in accordance with the Declaration of Helsinki and approved by the Ethics Committee of the Niigata University of Health and Welfare, Niigata, Japan (approval number 19170-231019). The participants were informed of data confidentiality and the study details, and provided written informed consent to participate in the study.

### Procedures

#### High-Intensity Exercise with a Progressive Load

The participants performed high-intensity exercises with progressive loading using an ergometer (828E; Monark, Sweden). They performed an ergometer exercise for 3 min at a load of 0.5 KP as a warm-up, followed by a 1-min rest period. After the break, males and females performed ergometer exercises with a progressive increase in the load of 0.5 KP/min. Participants performed the ergometer exercise and were instructed to pedal on time using a metronome (Smart Metronome; Tomohiro Ihara, Japan) set at 60 bpm. The exercise was stopped when the pedaling speed could no longer be sustained. During the exercise, strong verbal encouragement was provided to elicit maximal effort. Exercise time and heart rate (HR) pre-and post-intervention were measured during each session using a OH1 optical HR sensor (Polar, Finland) worn on the arm.

#### FR Intervention

Figure 1 shows the FR intervention method using a foam roller (Stretch Roll SR-002, Dream Factory, Umeda, Japan). A physical therapist instructed the participants prior to the intervention. FR intervention was initiated with knee flexors, followed by knee extensors. FR interventions were performed on the right and left sides. The FR area of the knee flexor muscle group was defined as the area proximal from the knee fossa to the sciatic tuberosity.^18^ The knee extensor was defined as the superior anterior iliac spine at the top of the patella.^19,20^ One FR cycle was defined as one distal rolling movement followed by one proximal rolling movement performed within 4 s.^21^ The intervention intensity was defined as the maximum intensity that the participants could tolerate (Kasahara et al. 2022). The intervention time for each area was 60 s, and the participants were instructed to prepare for the intervention for the next area during a 20-s interval. We used a metronome (Smart Metronome; Tomohiro Ihara, Japan) operating at 60 bpm for control.

**Figure 1.**
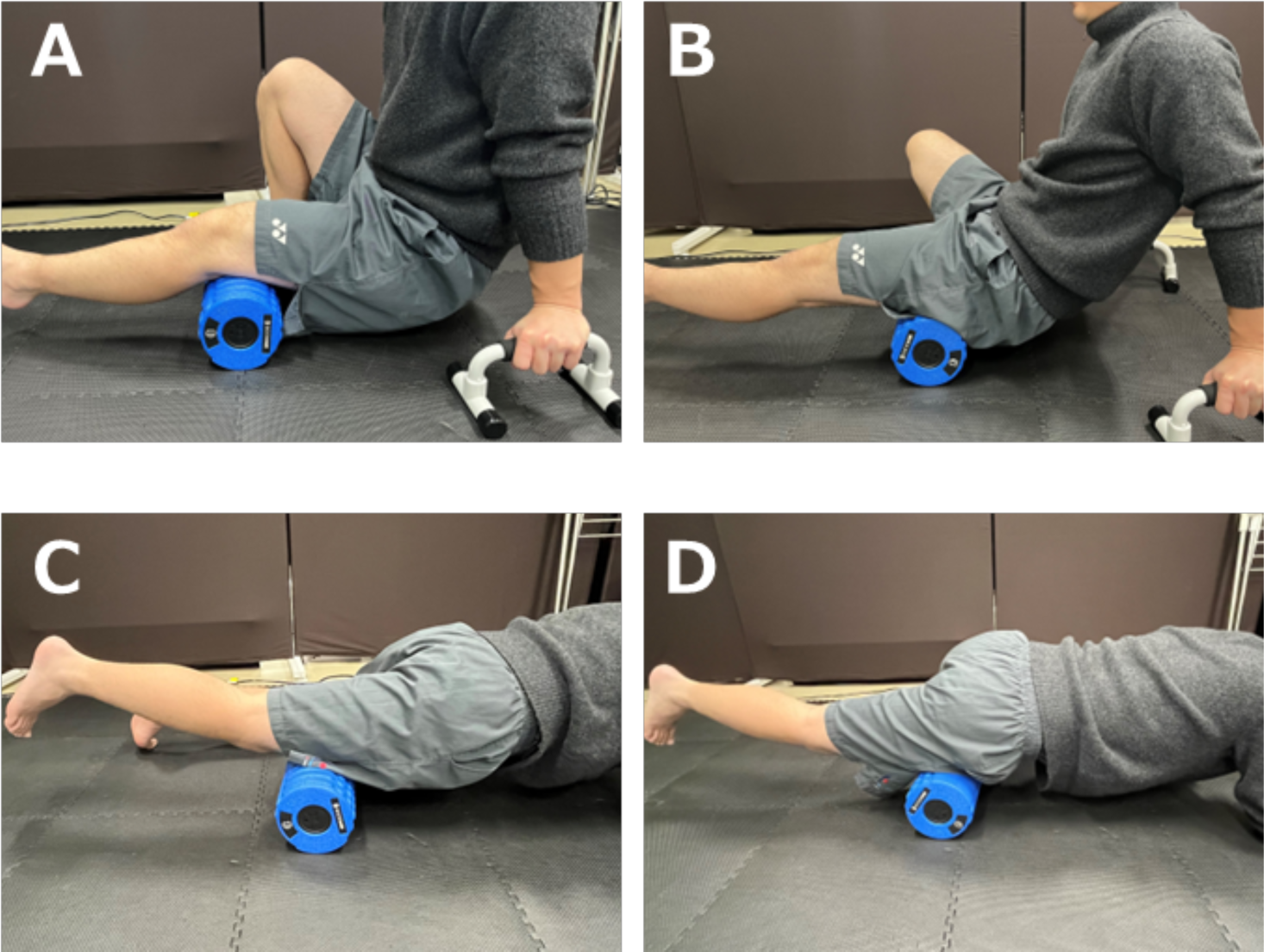
Intervention method for knee flexors and knee extensors. Panels A and B show the intervention method for the knee flexors. A push-up bar is used, with the sole on the non-intervention side in contact with the ground. A: shows the distal part of the knee flexors, and B: shows the proximal part. FR intervention is performed as shown from panel A to B. Panels C and D: the intervention for the knee extensors in which the non-intervention knee is instructed to touch the ground. C: the distal part of the knee extensors, and D: the proximal part. C to D: FR intervention is performed bilaterally for 60 s at a speed of 1 repetition every 2 s in the range specified for each muscle.

#### Psychological Measurements

POMS2 contains 35 items and evaluates seven mood states (anger-hostility, confusion-bewilderment, depression-ejection, fatigue-inertia, tension-anxiety, vigor-activity, and friendliness). This study used only a 10-item subset of vigor-activity and fatigue–inertia to reduce the psychological burden on the participants.

We recorded the rate of perceived exertion (RPE) before and after exercise to assess psychological exercise intensity in the whole body and lower extremities.

#### Blood Lactate Concentration

We used a Lactate Pro 2 (AKRAY, The Netherlands), a handheld point-of-care analyzer, for enzymatic amperometric detection. It requires 0.3 μl of a whole blood sample and a short time (15 s) to measure the lactate level. Lactate Pro 2 measurements range from 0.5 to 25.0 mmol/L. Therefore, only lactate values between 0.5 and 25 were included in this study.

#### Cognitive task performance

We employed Schneider’s spatial Stroop task to measure executive function.^22^ This task was created using web-building platforms (Lab.js v19.1.0).^23^ In each experiment, the participants made a spatially compatible key-press response to classify the spatial meaning of a stimulus presented at an irrelevant position. The stimulus meaning and position were either matched (congruent) or mismatched (incongruent) in the trial, with 16 congruent and 16 incongruent stimuli. Visual alert cues (squares) were briefly presented at two potential stimulus positions 500 ms before stimulus onset. Although previous studies used multiple stimulus types (arrows or words) and spatial dimensions (horizontal or vertical), this study employed arrow stimulus-type tasks. In addition, since preliminary experiments revealed that reaction times tended to be slower in vertical than in horizontal spatial dimension tasks, and the load on the task was assumed to be higher (fatigue was more likely to occur), vertical spatial dimension tasks were used in the present study.

Participants pressed a key indicating up (“y”) or down (“b”) to classify the direction of an up-or down-pointing arrow positioned above or below fixation (Figure 2). The correct answer rates assigned to yes and no were 50%. Each stimulus was separated by an inter-stimulus interval showing a fixation cross for 2 seconds to avoid prediction of the timing of the subsequent trial. The stimulus remained on the screen until the patient responded or for 1 s. This study adopted Stroop interference, a specifically defined cognitive process, to elucidate the effects of an acute bout of exercise on executive function by calculating the incongruent–congruent contrast, which was assumed to represent Stroop interference.

**Figure 2.**
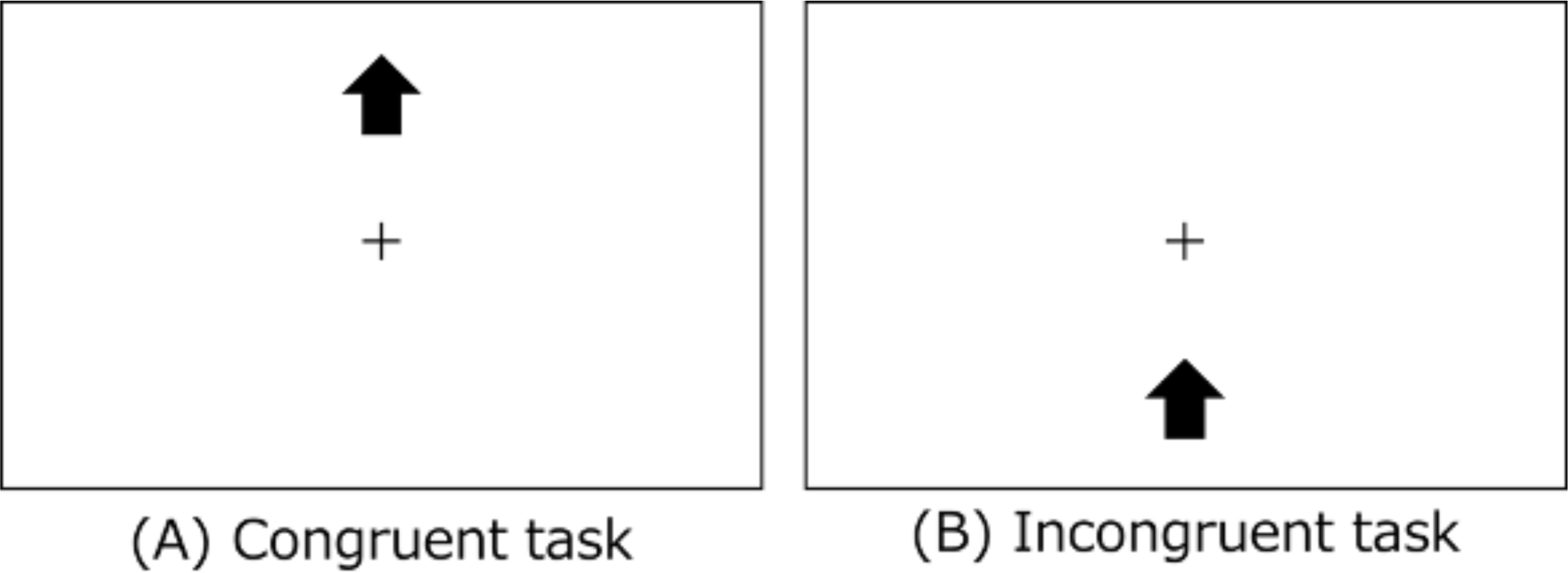
Example of Schneider’s spatial Stroop task. In the Congruent task (A), the displayed position and the direction of the arrow match as shown. In the Incongruent task (B), the position displayed on the screen and the direction of the arrow are mismatched, as shown.

Data from 16 participants (eight males and eight females) were used only for the cognitive task because data from two of the 18 participants could not be obtained.

### Statistical Analyses

R software (4.3.2), Rstudio (2023.12.0 + 369), “EZR on R Commander” (12), and the “anovaun” package were used for data analyses (The R Foundation, Vienna, Austria). A three-way analysis of variance (ANOVA) was performed to compare conditions, times, and sex. First, Mauchly’s sphericity test was used to determine whether sphericity was maintained. When this assumption was confirmed, a three-way ANOVA with Greenhouse–Geisser’s epsilon correction was performed; otherwise, only a three-way ANOVA was performed. If the 3-way ANOVA was not significant, a 2-way ANOVA was performed. The effect size classification was set where η*_p_*^2^ < 0.01 was considered small; 0.02–0.1, medium; and >0.1, large.^24^ A corresponding *t* test with Holm’s correction was used to determine significant differences obtained from the ANOVA. Pearson’s correlation analyses were performed to clarify the relationships between the parameters and executive performance. Statistical significance was set a priori at *P* < .05.

## RESULTS

### Physiological Parameters

Table 2 presents the results of the HR, RPE, total exercise time, and kp-max in the control and FR conditions. No significant differences in the pre-test values were observed between the two conditions for any of the measures. HR increased significantly (*P* < .01) immediately post-exercise in both conditions (control: 101.9±14.4, FR: 97.7±22.5). RPE (leg/body) was significantly (*P* < .01) higher at post-exercise (leg, control: 10.4±3.0, FR: 10.6±2.9; body, control: 9.1±2.6, FR: 8.9±3.5) and 5 min into the resting period (leg, control: 6.7±3.8, FR: 6.2±3.1; body, control: 5.1±3.9, FR: 4.7±3.3) as compared to pre-exercise in both conditions but significantly (*P* < 0.01) lower at 5 min into the resting period when compared with post-exercise (leg, control: −3.7±2.0, FR: −4.4±2.4; body, control: −4.0±2.3, FR: −4.2±3.0). In addition, sex differences were observed in the total exercise time and post-exercise load, which were significantly (*P* < .01) higher in males than in females. However, no significant differences were observed between the two conditions.

**Table 2.** Physiological parameters of each condition before (pre), immediately after exercise (post), and after intervention (5 min) for each sex (mean±SD)

| Conditions and variables | Males | Females |
| --- | --- | --- |
| Control |  |  |
| HR pre (bpm) | 76.9±10.4 | 75.5±6.4 |
| HR post (bpm) | 181.3±11.9* | 174.4±12.8* |
| RPE pre: leg / body (point) | 6.4±0.7 / 6.5±0.7 | 7.9±2.2 / 8.1±2.3 |
| RPE post: leg / body (point) | 18.3±1.5* / 16.8±2.1* | 16.4±1.8* / 15.8±1.6* |
| RPE 5 min: leg / body (point) | 14.7±2.9*† / 13.0±3.4*† | 12.5±2.2*† / 11.5±2.7*† |
| Total exercise time (sec) | 651.0±87.0 | 405±92.5** |
| kp max (a.u.) | 3.4±0.4 | 2.3±0.4** |
| Foam Rolling |  |  |
| HR pre (bpm) | 77.1±12.2 | 81.6±6.6 |
| HR post (bpm) | 181.6±16.2* | 170.9±25.2* |
| RPE pre: leg / body (point) | 6.5±1.0 / 6.6±1.0 | 7.8±2.2 / 8.4±2.3 |
| RPE post: leg / body (point) | 18.6±1.4*/ 17.2±2.6* | 16.5±2.0*/ 15.1±2.7* |
| RPE 5 min: leg / body (point) | 14.4±2.4*† / 12.7±3.2*† | 11.9±2.0*† / 11.4±2.1*† |
| Total exercise time (sec) | 615.5±138.5 | 396.6±91.7** |
| kp max (a.u.) | 3.2±0.6 | 2.3±0.4** |

\* Significant difference in pre-exercise conditions ( $P < 0.01$ ); † significant difference in post-exercise conditions ( $P < .01$ ); \*\* significant difference in males ( $P < .01$ ). HR, heart rate; RPE, rating of perceived exertion; SD, standard deviation.

Figure 3 presents the changes in lactate levels. A three-way ANOVA (conditions vs. time vs. sex) showed no significant interactions (*F* (2, 32) = 1.0047, *P* = 0.3774, η*_p_*^2^ = 0.0591). A two-way ANOVA (conditions vs. time) showed a significant interaction (*F* (2,32) = 4.0433, *P* < .05, η*_p_*^2^ = 0.2017) and a main effect of time (*F* (2, 32) = 275.1504, *P* < .0001, η*_p_*^2^ = 0.9450). The post-hoc test results in the control condition indicated significantly higher values at post-exercise (10.6±2.0 mmol/L) and 5 min into the resting period (9.6±2.9 mmol/L) than those at pre-exercise (*P* < .01), although no significant difference was observed between post-exercise and 5 min into the resting period (−1.0±2.1 mmol/L). In the FR condition, post-exercise (16.3±2.0 mmol/L) and 5 min into the resting period (9.0±2.5 mmol/L) values were significantly higher than pre-exercise (*P* < .01). In addition, results at 5 min into the resting period (−7.3±3.0 mmol/L) were significantly lower than that at post-exercise (*P* < .01).

**Figure 3.**
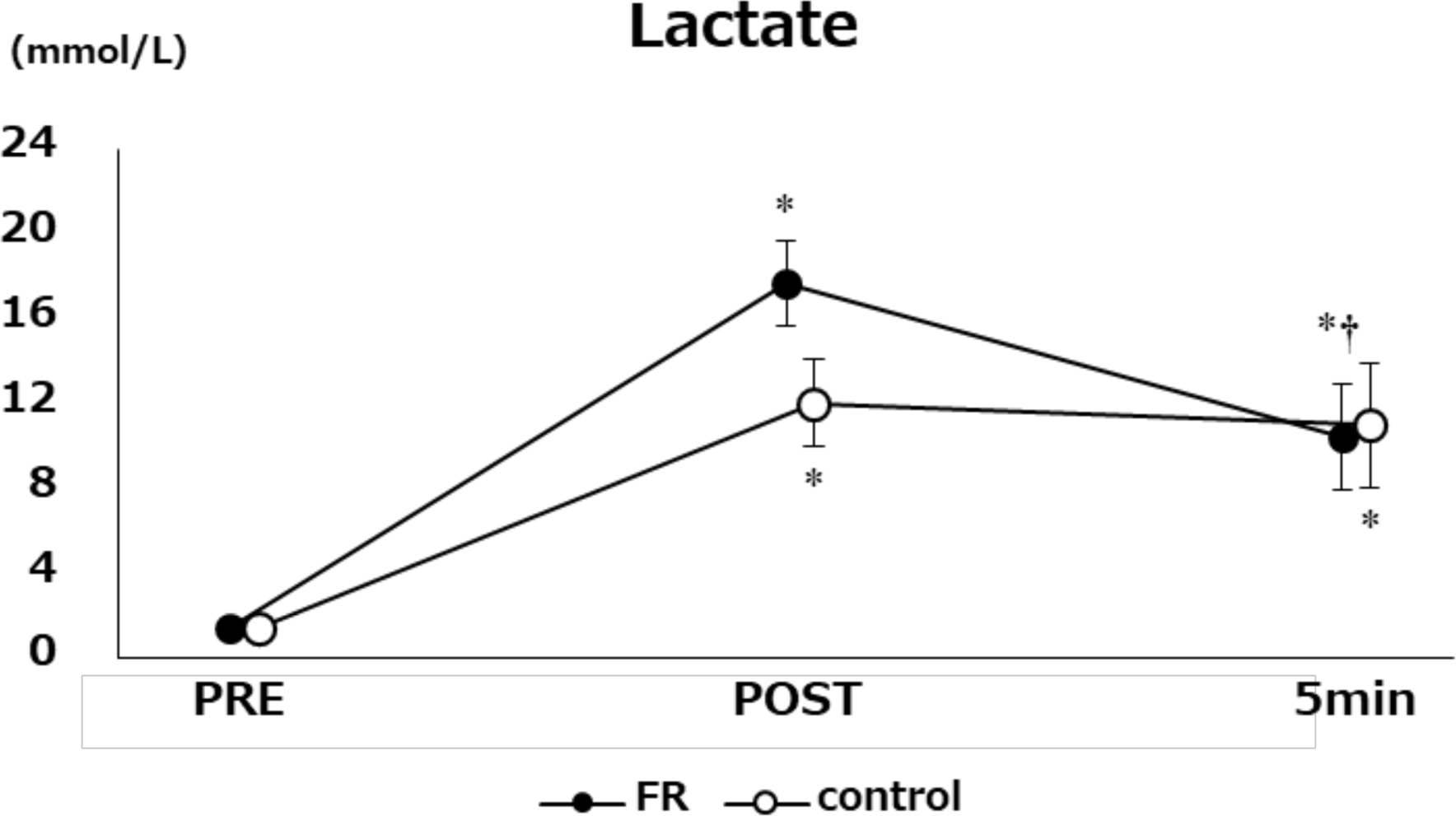
Changes in lactate at pre, post, and 5 min after each condition. *: Significant difference in pre-exercise conditions (*P* < .01); †: Significant difference in post-exercise conditions (*P* < .01). FR, foam rolling.

### Psychological Parameters

A three-way ANOVA (conditions vs. time vs. sex) showed no significant interaction between vigor (*F*(2, 32) = 1.6850, *P* = .2015, η*_p_*^2^ = 0.0953) and fatigue (*F*(2, 32) = 1.0022, *P* = .3783, η*_p_*^2^ = 0.0589) in POMS2. A two-way ANOVA (conditions vs. time) showed a main effect of time on vigor (*F*(2, 32) = 7.5235, *P* < .01, η*_p_*^2^ = 0.3198) and fatigue (*F*(2, 32) = 163.5833, *P* < .0001, η*_p_*^2^ = 0.9109). A post-hoc test using Holm correction revealed that vigor was significantly lower in post-exercise (control: −2.2±4.6, FR:-4.1±5.4) than in pre-exercise conditions (*P* < .01). Fatigue was significantly higher at post-exercise (control: 12.7±4.1, FR: 12.4±3.7) and 5 min into the resting period (control: 8.1±4.4, FR: 8.4±4.8) as compared to that at pre-exercise (*P* < .0001). In addition, post-exercise values (control: −4.6±2.9, FR: −4.0±3.2) were significantly higher than those at 5 min into the resting period (*P* < .0001).

### Relationship Between Parameters and Performance

Figure 4 displays the changes in lactate levels and executive function pre-and post-intervention. A significant correlation was observed for the FR condition (*r* = 0.41, *P* = .086). Sex-specific results showed a significant correlation for males (*r* = 0.706, *P* < .05), but not for females (*r* = 0.13, *P* = .747). No significant correlation was found in the control group (*r* = −0.38, *P* = .118).

**Figure 4.**
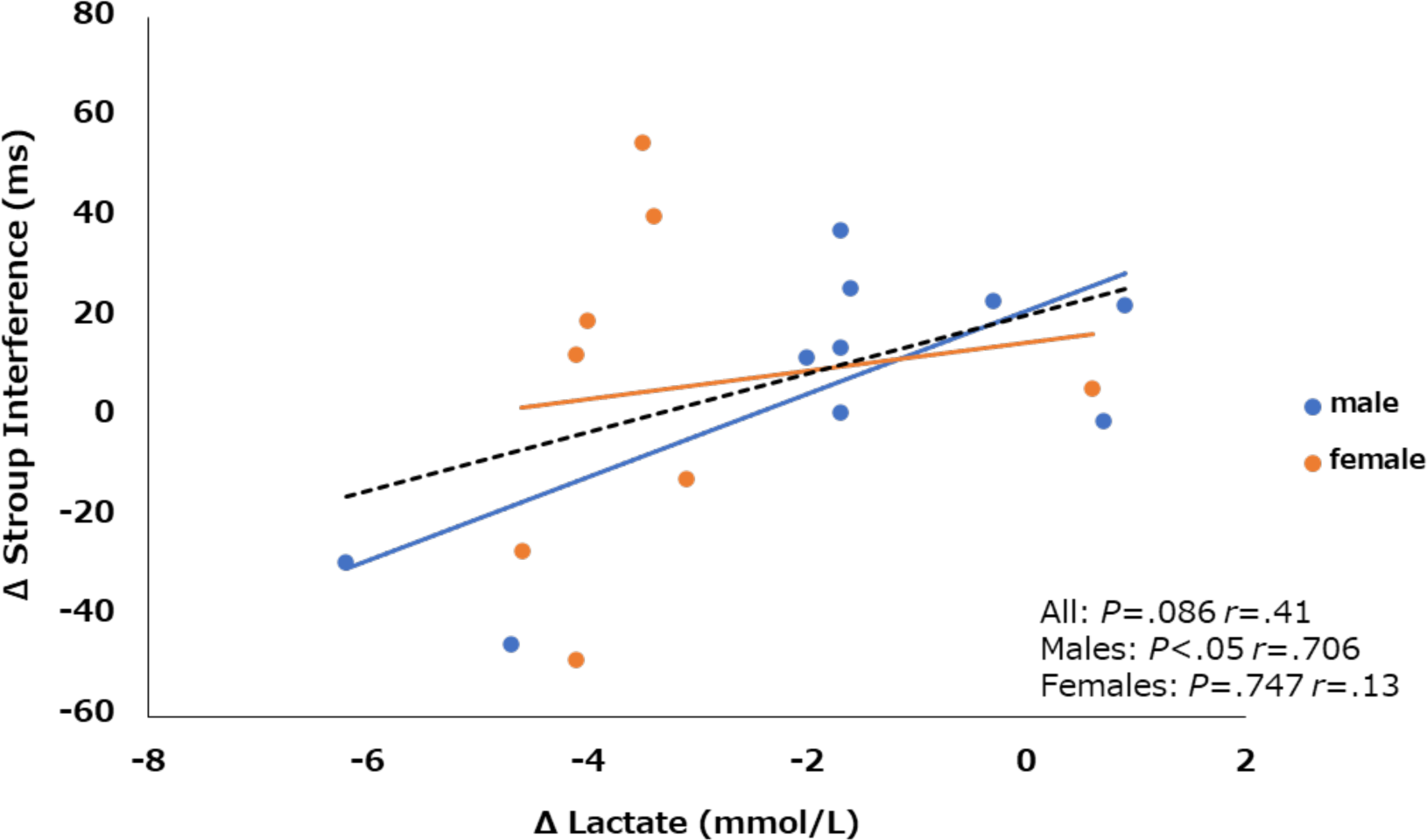
Correlation between the changes in lactate and Stroop interference in the FR condition. The dotted lines indicate the regression lines for sexes; blue for males and orange for females. FR, foam rolling.

Figure 5 presents the changes in lactate levels and RPE (leg/body) before and after the intervention. No significant correlation was found in the FR condition; however, a significant correlation was found in the control condition (leg: *r* = 0.778, *P* < .01; body: *r* = 0.669, *P* < .01).

**Figure 5.**
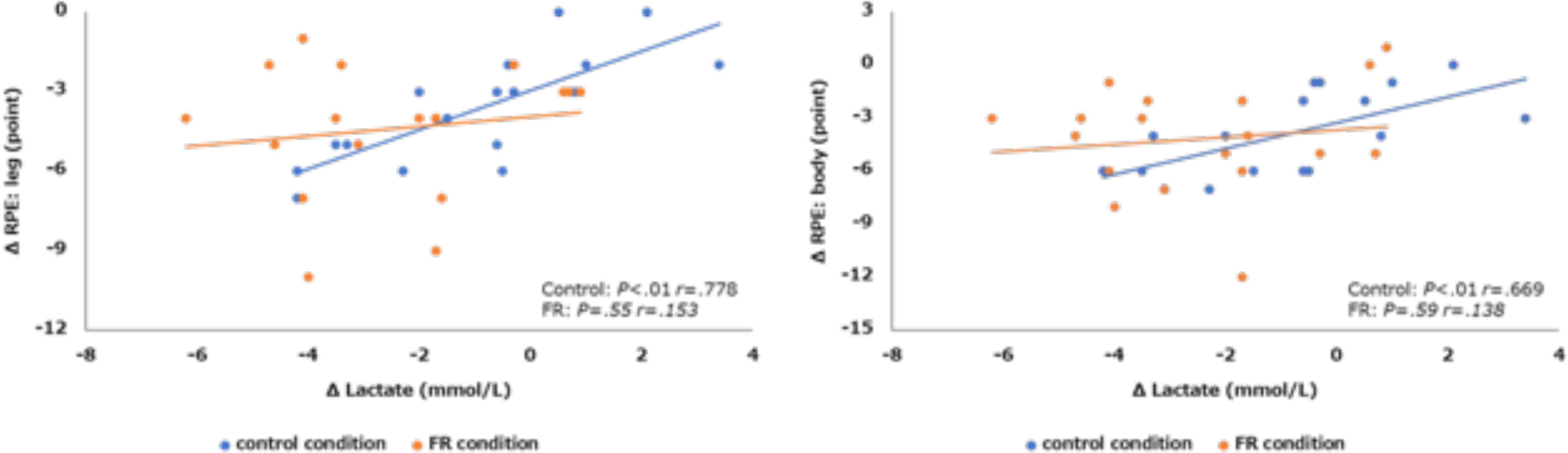
Correlations between the change in lactate for the RPE leg and whole body in FR and control conditions. FR, foam rolling; RPE, rating of perceived exertion.

## DISCUSSION

We examined the acute effects of FR intervention on lactate clearance, psychological parameters, and cognitive function after a progressively increasing load exercise, resulting in exhaustion to propose FR as a new recovery method that athletes can implement in a limited space. The results revealed that arterial blood lactate concentration, which increased after exhaustive exercise, decreased in the FR condition regardless of sex, indicating that FR intervention enhanced lactate clearance. Interestingly, cognitive function improved after FR intervention as compared to those with greater lactate clearance, and this effect was significantly greater in males than in females. This finding indicates that FR intervention may be useful in improving exercise-induced cognitive decline, particularly in males.

This study showed significant increases in HR at pre-and post-exercise in both the FR and control conditions, without significant differences between them. In addition, the Karvonen method^25^ demonstrated that exercise intensity at post-exercise was 85.3 ± 11.0% in the FR condition and 83.6% ± 10.4% in the control condition, with no significant difference between the two conditions. Based on these results, we concluded that all participants performed the same amount of exercise under both conditions.

First, we examined the effects of FR intervention on lactate clearance. In the present study, ergometric exercise was performed with progressive loading by applying a strong load to the lower extremities, and the FR intervention targeted the bilateral knee extensors and flexors. The arterial blood lactate concentrations increased significantly immediately after exercise under both conditions, and no difference was observed between the conditions. Thereafter, no lactate concentration changes were observed in the control condition. However, in the FR condition, the arterial blood lactate concentration significantly decreased post-intervention. These results indicated that FR intervention immediately after exhaustive exercise was useful in increasing lactate clearance. A previous study reported that increased arterial blood lactate concentrations after high-intensity exercise were significantly lower in active recovery groups that performed low-or high-intensity exercises than in passive recovery groups that did not receive any intervention.^26,27^ Interestingly, no significant difference in the increasing effect of the ROM was observed between the FR and sham rolling conditions, which simulated FR intervention movements.^28^ These findings demonstrated that active recovery, such as FR, may increase blood flow in active tissues and lactate clearance.^29^ In addition, FR intervention increases regional perfusion^30^ ) and, therefore, may further increase lactate clearance.

Second, the effects of the FR intervention on executive function were examined. In the present study, executive function task performance did not decrease before or after exercise until exhaustion, and progressive-load exercises leading to exhaustion did not induce cognitive fatigue. Previous study showed progressive exercise to exhaustion caused impaired cognitive function, including executive function^31^; however, its effects are not always clear. Whether laboratory training impairs executive function varies greatly from person to person, and further testing is required. Exercise in extreme environments, such as hypoxic environments, decreases cognitive function.^2,3^ To verify whether FR intervention is effective in restoring exercise-induced cognitive decline, it should be investigated under different exercise conditions such as a hypoxic environment.

This study demonstrated that the amount of lactate change was significantly correlated with changes in executive function task performance immediately after exercise and post-FR intervention in the FR condition. This result indicates that executive function may recover in patients with increased lactate clearance. Previous studies have reported that elevated blood lactate levels after exhaustive exercise are associated with significantly worsened attentional processes.^13^ Therefore, increased lactate clearance due to FR may contribute to improved cognitive performance due to increased lactate levels. Although we did not observe a decline in executive function in this study, it may be possible to propose a new effect of FR by testing it under experimental conditions in which a decline in executive function occurs.

We examined the effects of the FR intervention on psychological parameters. The results indicated that FR intervention did not affect the vigor or fatigue of POMS2 and support the findings of a previous study^32^ in which FR intervention was administered to female basketball players and had no effect on perceived recovery (total quality recovery) or fatigue level. However, a larger effect size in professional male soccer players (Cohen’s d = 1.08) was reported for total perceived recovery in the FR intervention condition than in the control condition after 24 h of training.^33^ These differences in results may be due to the FR intervention methods. Pernigoni et al. (2023) and Rey et al.^33^ performed a 20-min FR intervention on the entire lower extremity, including the quadriceps, hamstrings, triceps, as well as abductor and adductor muscle groups. Conversely, this study found that the FR intervention lasted 4 min for knee extensors and knee flexors. A capacitance–response relationship has been found for the effect of FR on the ROM increase.^34^ The enhancement of psychological parameters through FR may also be associated with the capacity–response relationship, necessitating further research for a clearer understanding.

Regarding RPE, which represents perceived exertion, the results revealed a greater decrease in lactate in the control condition was associated with a lower RPE. However, in the FR condition, no significant correlation was observed between the changes in lactate and RPE. These differences may be related to the FR intervention methods. A push-up bar was used as the FR intervention method for the knee flexors (Figure 1A, B). Furthermore, participants performed the intervention for the knee extensors in a plank posture (Figure 1C, D). In this study, since we examined the intervention effects in participants who did not use a foam roller daily, the FR intervention itself may have affected the RPE. Therefore, although the lactate clearance increased in the FR condition, the effect of improved perceived exertion might not have occurred. The on-field implementation of FR interventions requires careful consideration of whether priority should be given to the recovery of physiological parameters or subjective fatigue levels.

This study has a few limitations. First, the mechanisms underlying these results remain unknown. The results indicated that FR intervention increased lactate clearance and concomitantly affected executive function. However, the detailed mechanism by which this occurs remains unclear. Future studies should examine the effects of FR intervention on regional and cerebral perfusion and changes in lactate clearance and cognitive function. Second limitation is the target population. As mentioned above, FR intervention for male athletes showed fatigue recovery effects,^33^ but not for female basketball players .^32^ Therefore, differences between those with exercise habits and older adults, as well as differences in athletic specialties and sex among athletes, should be examined in the future. Finally, the effect of FR intervention on exercise-induced cognitive decline (cognitive fatigue) was discussed.

## PRACTICAL APPLICATION

FR intervention may be a useful recovery method to improve muscle function and reduce the decrease in cognitive performance. However, the FR intervention was not effective in restoring RPE. Athletes and coaches should carefully consider whether the recovery of physiological parameters or RPE should be prioritized when implementing FR interventions in the field.

## CONCLUSIONS

The results of the present study revealed that the acute effect of FR intervention after exhaustive exercise in healthy young adults was an increase in lactate clearance. Furthermore, the greater the effect, the greater the executive function recovery. However, FR intervention did not decrease RPE after exhaustive exercise. Although this study was conducted in young adults without exercise habits, further validation in athletes will allow us to propose a useful FR intervention program.

## ACKNOWLEDGMENTS

The authors gratefully acknowledge all the participants involved in this study.

## Data availability

All data generated or analyzed during this study are included in this article.

